# Increased complement activation is a distinctive feature of severe SARS-CoV-2 infection

**DOI:** 10.1101/2021.02.22.432177

**Authors:** Lina Ma, Sanjaya K. Sahu, Marlene Cano, Vasanthan Kuppuswamy, Jamal Bajwa, Ja’Nia McPhatter, Alexander Pine, Matthew Meizlish, George Goshua, C-Hong Chang, Hanming Zhang, Christina Price, Parveen Bahel, Henry Rinder, Tingting Lei, Aaron Day, Daniel Reynolds, Xiaobo Wu, Rebecca Schriefer, Adriana M. Rauseo, Charles W. Goss, Jane A. O’Halloran, Rachel M. Presti, Alfred H. Kim, Andrew E. Gelman, Charles Dela Cruz, Alfred I. Lee, Phillip Mudd, Hyung J. Chun, John P. Atkinson, Hrishikesh S. Kulkarni

**Author notes:** **CORRESPONDING AUTHOR**: Hrishikesh S. Kulkarni, MD, MSCI, Division of Pulmonary and Critical Care Medicine, Washington University School of Medicine, 660 S Euclid Avenue, Campus Box 8052, St. Louis, MO 63110, United States.

## Abstract

Complement activation has been implicated in the pathogenesis of severe SARS-CoV-2 infection. However, it remains to be determined whether increased complement activation is a broad indicator of critical illness (and thus, no different in COVID-19). It is also unclear which pathways are contributing to complement activation in COVID-19, and, if complement activation is associated with certain features of severe SARS-CoV-2 infection, such as endothelial injury and hypercoagulability. To address these questions, we investigated complement activation in the plasma from patients with COVID-19 prospectively enrolled at two tertiary care centers. We compared our patients to two non-COVID cohorts: (a) patients hospitalized with influenza, and (b) patients admitted to the intensive care unit (ICU) with acute respiratory failure requiring invasive mechanical ventilation (IMV). We demonstrate that circulating markers of complement activation (i.e., sC5b-9) are elevated in patients with COVID-19 compared to those with influenza and to patients with non-COVID-19 respiratory failure. Further, the results facilitate distinguishing those who are at higher risk of worse outcomes such as requiring ICU admission, or IMV. Moreover, the results indicate enhanced activation of the alternative complement pathway is most prevalent in patients with severe COVID-19 and is associated with markers of endothelial injury (i.e., Ang2) as well as hypercoagulability (i.e., thrombomodulin and von Willebrand factor). Our findings identify complement activation to be a distinctive feature of COVID-19, and provide specific targets that may be utilized for risk prognostication, drug discovery and personalized clinical trials.

**SUMMMARY:** Complement has been implicated in COVID-19. However, whether this is distinctive of COVID-19 remains unanswered. Ma et al report increased complement activation in COVID-19 compared to influenza and non-COVID respiratory failure, and demonstrate alternative pathway activation as a key marker of multiorgan failure and death.

## INTRODUCTION

Morbidity and mortality associated with SARS-CoV2 infection (i.e., COVID-19) have been attributed to a hyperinflammatory phase (Vabret *et al*., 2020; Zhou *et al*., 2020; Wu *et al*., 2020; Blanco-Melo *et al*., 2020). Specifically, approximately 7-10 days after clinical onset, a subset of patients require hospitalization, ICU admission, and mechanical ventilation, and may ultimately die due to their illness (Wiersinga *et al*., 2020). However, the components of the immune response that contribute to critical illness in COVID-19 remain incompletely understood. For example, although certain cytokines such as IL-6, G-CSF, IL-1RA, and MCP-1 predict death in COVID-19, their circulating levels are no different when measured in patients with other viral infections, such as influenza (Mudd *et al*., 2020). However, the clinical presentation and autopsy findings of patients with COVID-19 indicate that in at least some of these patients, there may be a distinct immunological response, which is responsible for the mortality rate in excess of other viral illnesses (i.e., influenza) (Cobb *et al*., 2020; Xie *et al*., 2020), and results in certain coagulopathic events such as microscopic and macroscopic thrombi occurring more commonly in COVID-19 (Ackermann *et al*., 2020; Bradley *et al*., 2020; Carsana *et al*., 2020; Fox *et al*., 2020; Mei *et al*., 2020). Understanding the underlying mechanisms of these relatively unique aspects of COVID-19 is crucial for targeting therapies, and, may provide insights into the pathogenesis of acute respiratory distress syndrome on a broader scale.

The complement system, one of the first lines of the host defense and a key player in the innate immune response, has been implicated in the pathogenesis of severe COVID-19 (Holter *et al*., 2020; Java *et al*., 2020; Skendros *et al*., 2020). Features of disease such as hypercoagulability and tissue necrosis, as well as genetic factors have increased the suspicion that the complement system contributes to severe illness (Java *et al*., 2020; Perico *et al*., 2020; Ramlall *et al*., 2020; Valenti *et al*., 2021). The system can be activated by three arms – the classical, lectin or the alternative pathway (Kulkarni *et al*., 2018). Although a prevailing hypothesis is that the N-protein of coronaviruses triggers MASP2-mediated complement activation and thus drives disease severity via the lectin pathway (Gao *et al*., 2020), in vitro studies have suggested that spike proteins (subunits S1 and S2) of SARS-CoV2 activate the alternative pathway (Yu *et al*., 2020). Regardless of the initial activation step, the system converges on the cleavage of C3 and subsequently C5 to anaphylatoxins that facilitate vasodilation, chemotaxis and thrombosis (C3a, C5a). Further, activation of the system facilitates opsonization (via C3b), and membrane attack complex formation (MAC, i.e., C5b-9) (Kulkarni *et al*., 2018). Accordingly, multiple studies have demonstrated elevation of C5a and sC5b-9 in patients with COVID-19 (Carvelli *et al*., 2020; Cugno *et al*., 2020; Holter *et al*., 2020), as well as deposition of activated complement proteins in injured tissues and organs (Magro *et al*., 2020; Macor *et al*., 2021). As a result, these studies have created a precedent for targeting the complement system in multiple ongoing phase II and phase III clinical trials employing complement inhibitors in COVID-19 (Declercq *et al*., 2020; Mastellos *et al*., 2020; Smith *et al*., 2020; Vlaar *et al*., 2020; Mansour *et al*., 2021).

Most studies addressing the role of complement in COVID-19 have not included acute respiratory infection cohorts without COVID-19. Therefore, it is unclear whether complement activation is unique to severe COVID-19, or simply a broader feature of critical illness. Additionally, which arms of the complement cascade contribute to complement activation in patients with COVID-19, remains to be defined. Finally, whether complement activation in vivo is associated with certain distinctive features of COVID-19 (i.e., endothelial injury and hypercoagulability) is also unclear. Here we report that increased complement activation is an immunological feature of COVID-19, which distinguishes those developing severe illness. Using two independent cohorts, we identify components of the alternative pathway are markedly elevated in patients with severe COVID-19. Our findings may potentially refine the approach for therapeutically targeting the complement system in severe SARS-CoV2-infection.

## RESULTS

### Markers of complement activation are higher in COVID-19 compared to non-COVID-19 respiratory failure

We first sought to assess complement activation in patients hospitalized with COVID-19, versus those with non-COVID-19-related illness. We compared plasma sC5b-9 levels in 134 patients with COVID-19 at WUSM (**Table 1, Figure 1A)** — with two independent cohorts of non-COVID-19 acute respiratory illnesses (**Table 1, Figures S1A and S1B**). The first comparison was with the EDFLU cohort of 54 patients presenting with influenza at WUSM. Patients hospitalized with COVID-19 had significantly higher median plasma sC5b-9 levels [666.3 (interquartile range, 429.7 – 980.1) ng/mL] compared to those with influenza [254.5 (154.5 – 403.8) ng/mL, p<0.0001, **Figure 1B**]. Given that a minority of the influenza cohort required invasive mechanical ventilation (IMV, 6/54, 11%), we also compared the plasma sC5b-9 levels of patients in our COVID-19 cohort with patients in our IPS cohort, all of whom required IMV in the ICU for non-COVID-19-related acute respiratory failure (n=22). Plasma sC5b-9 levels were higher in patients with COVID-19 when compared to the IPS cohort [243.5 (95.62 – 352.1) ng/mL, p<0.0001, **Figure 1C**]. Plasma sC5b-9 levels in the COVID-19 cohort remained higher than the levels in the IPS cohort despite restricting the COVID-19 cohort to those admitted to the ICU (**Figure S1C)** and among those requiring IMV (**Figure S1D**). These observations suggested that patients with COVID-19 appear to have higher circulating markers of complement activation compared to patients with non-COVID-19-related acute respiratory infection. When restricted to those who died, patients in the COVID-19 cohort had higher plasma sC5b-9 levels [751.7 (575.2 – 1118) ng/mL, n=31] compared to the non-COVID-19 IPS cohort [173.3 (78.99 – 353.3) ng/mL, n=8, p < 0.0001, **Figure 1D**].

**Table 1.** Demographic characteristics of the cohorts.

|  | COVID-19 (n=134) | EDFLU (n=54) | ISF (n=22) |
| --- | --- | --- | --- |
| **Demographics** |  |  |  |
| Age in years, median (IQR) | 65 (55.5 – 72.9) | 53.5 (43 – 63.5) | 58 (36 – 69) |
| **Gender** |  |  |  |
| Female | 41% (55) | 50% (27) | 50% (11) |
| Male | 59% (79) | 50% (27) | 50% (11) |
| **Clinical Characteristics** |  |  |  |
| Hospital Admission | 92.5% (124) | 96.3% (52) | 100.0% (22) |
| ICU Admission | 53.7% (72) | 24.1% (13) | 100.0% (22) |
| Invasive Mechanical Ventilation | 21.6% (29) | 11.1% (6) | 100.0% (22) |
| In-Hospital Mortality | 22.4% (30) | 3.7% (2) | 36.4% (8) |
| **Comorbidities** |  |  |  |
| Smoking History <sup>a</sup> | 46.3% (62) |  | 63.6% (14) |
| Chronic Lung Disease | 19.4% (26) | 22.2% (12) | 13.6% (3) |
| End-stage renal disease (ESRD)<sup>b</sup> | 6% (8) | 7.4% (4) | 4.5% (1) |
| Diabetes mellitus, type 2 | 52.2% (70) | 35.2% (19) | 18.2% (4) |

Abbreviations: COVID-19: patients presenting to the emergency department with SARS-CoV2 infection; EDFLU: patients presenting to the emergency department with influenza; ICU: intensive care unit; ISF: Immunity in Pneumonia and Sepsis (patients in the intensive care unit for non-COVID acute respiratory failure); IQR: interquartile range
<sup>a</sup> current or former smoker
<sup>b</sup>based on chronic dialysis requirement

**Figure 1.**
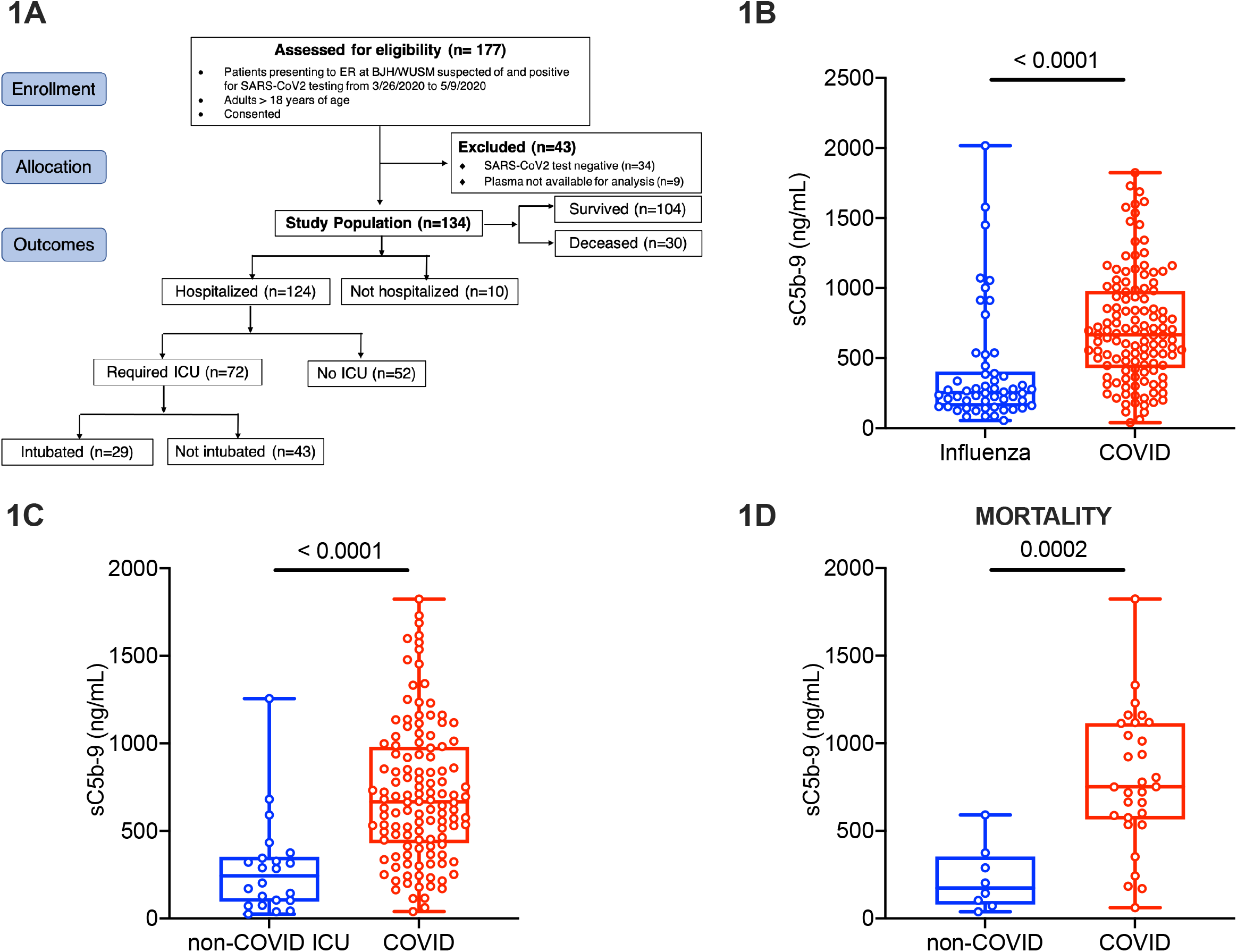
Markers of complement activation are unique to COVID-19 compared to non-COVID-19 respiratory failure. Plasma for determination of circulating markers of complement activation was obtained in patients with COVID-19 and influenza at Barnes-Jewish Hospital (BJH)/Washington University School of Medicine (WUSM). (A) CONSORT flow diagram showing patient enrollment, allocation and outcomes in the COVID-19 cohort. The CONSORT diagram for the influenza and non-COVID acute respiratory failure cohorts are in Figure S1. Box and whiskers plots of differences in sC5b-9 between (B) the influenza (EDFLU) and COVID-19 cohorts, (C) the non-COVID acute respiratory failure (Immunity in Pneumonia and Sepsis, IPS) and the COVID-19 cohorts, and (D) restricting the cohorts in Fig.1C to those who died. The center of the box represents the median value, and the length of the box represents the interquartile range. The whiskers represent the minimum and maximum values in each group. Statistical significance is determined using Mann-Whitney U test.

### Complement activation is associated with worse outcomes in two independent COVID-19 cohorts

Plasma sC5b-9 levels were higher in patients belonging to the WUSM COVID-19 cohort who required hospitalization [666.3 (429.7 – 980.1) ng/mL, n=124], compared to those who were discharged from the emergency room [326.2 (211.6 – 584.4) ng/mL, n=10, p=0.0097, **Figure S2A**], as well as in those requiring ICU admission (**Table 2, Figure 2A**). Plasma sC5b-9 levels were higher in patients who required IMV [922.8 (545.0 – 1198.0) ng/mL, n=29] compared to those who did not [600.8 (349.2 – 838.8) ng/mL, n=105, p=0.0034, **Figure 2B**]. This comparison held even when restricting the COVID-19 cohort to those who were hospitalized (**Figure S2B**), and those admitted to the ICU (**Figure S2C**). Patients with COVID-19 who died had higher plasma sC5b-9 levels [751.4 (565.2 – 1115) ng/mL, n=30] compared to those who survived the index hospitalization, although this did not reach statistical significance [600.0 (349.9 – 858.5) ng/mL, n=104, p=0.0666, **Figure 2C**]. We also measured sC5a, which is a product of C5 cleavage similar to C5b (that contributes to the formation of membrane attack complex, C5b-9). In the WUSM cohort, plasma sC5a correlated with sC5b-9 (ρ=0.4909, **Figure 2D**), and sC5a levels were significantly higher in patients with COVID-19 requiring ICU admission compared to those who did not (**Table 2, Figure S2D**).

**Table 2.**
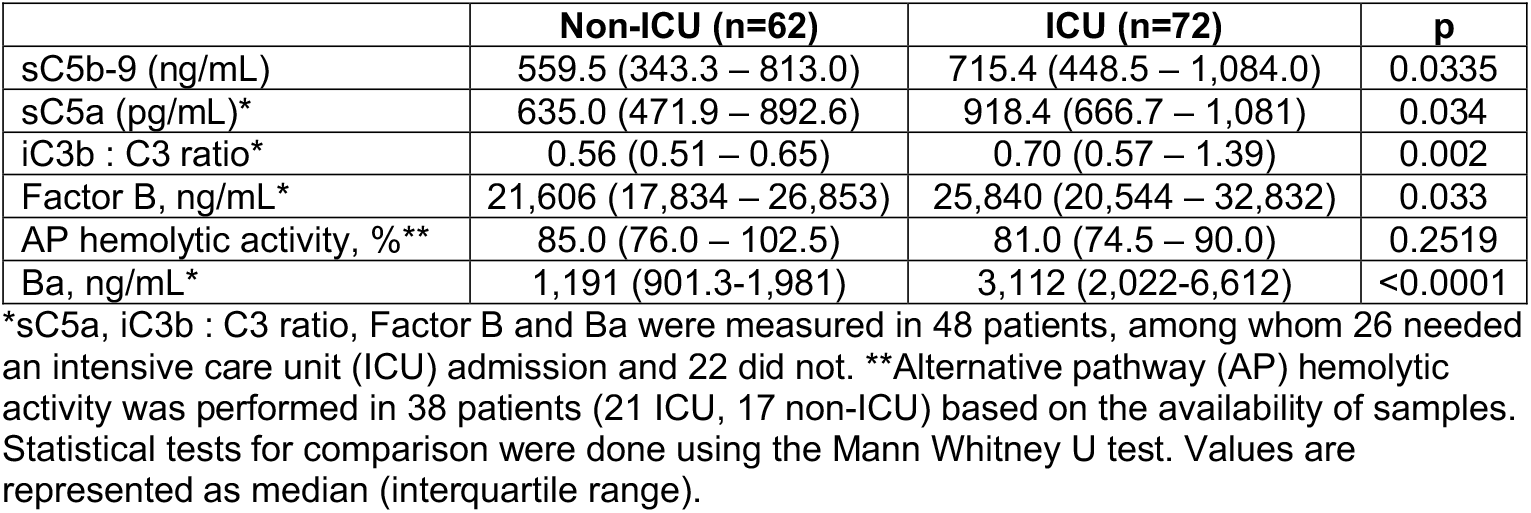
Complement analytes in the Washington University School of Medicine (WUSM) COVID-19 cohort.

**Figure 2.**
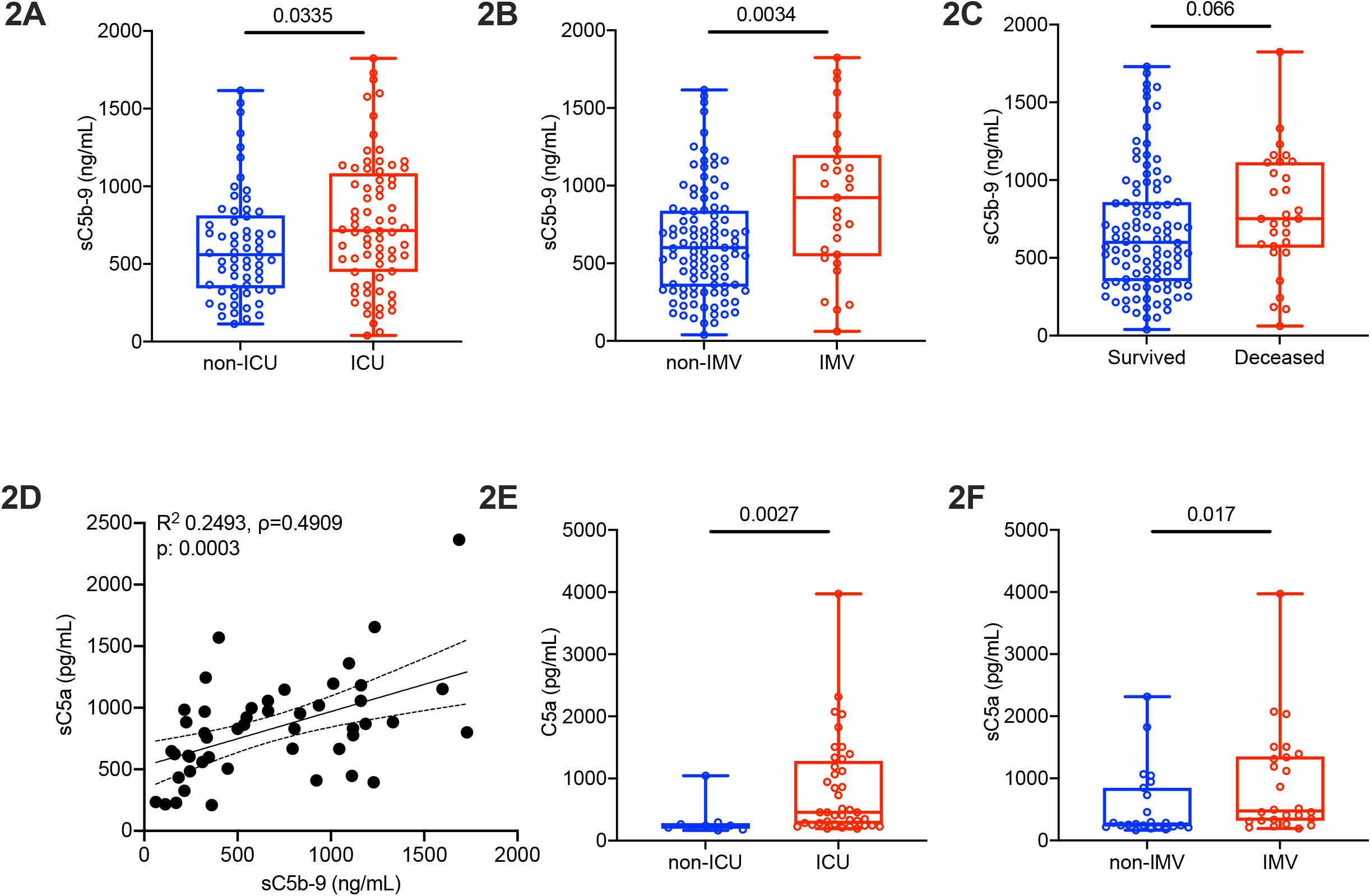
Complement activation is associated with worse outcomes in COVID-19 in two independent cohorts. Markers of complement activation were quantified in the plasma at WUSM and Yale University School of Medicine (Yale). Box and whiskers plots of sC5b-9 levels in the WUSM COVID-19 cohort in (A) patients requiring ICU admission versus those who did not, (B) patients requiring invasive mechanical ventilation (IMV) versus those who did not, and (C) patients who died versus those who survived. (D) A linear regression line shows the relationship between plasma levels of sC5b-9 and sC5a. The spline chart demonstrates the mean with 95% confidence intervals. R^2^ represents the goodness-of-fit. The degree of correlation is assessed using Spearman’s Rank Correlation Coefficient test (ρ=0.4909, 95% CI 0.2321 – 0.6848, n=48). In the Yale longitudinal cohort, concurrently measured sC5a levels are utilized to compare (E) patients requiring ICU admission versus those who did not, and (F) patients requiring IMV versus those who did not. The center of the box represents the median value, and the length of the box represents the interquartile range. The whiskers represent the minimum and maximum values in each group. Statistical significance is determined using Mann-Whitney U test.

To test whether our findings hold true at a center that had independently measured inflammatory markers in a similar time frame of patient enrollment, we utilized a second cohort from Yale School of Medicine, wherein sC5a had been prospectively measured in the plasma of patients hospitalized with COVID-19 (Yale cross-sectional cohort, n=23). within the first 24 h of hospital admission. In this cohort, plasma sC5a levels were significantly higher in those patients who requiring ICU admission (**Table 3, Figure S2E**). In this cohort, there were not enough patients to make a meaningful comparison regarding the need for IMV (n=2, 9%). Hence, we expanded the cohort to include those patients who had their first plasma sampled beyond the first day of hospital admission (Yale longitudinal cohort, n=49). Even in this expanded cohort, plasma sC5a levels remained significantly higher in hospitalized patients with COVID-19 requiring ICU admission [456.9 (269.2 – 1282) pg/mL, n=40] versus those who did not [243.9 (193.3 – 280.3) pg/mL, n=9, p=0.0027, **Figure 2E**]. Additionally, among those patients with COVID-19 who were hospitalized, plasma sC5a levels were significantly higher in those requiring IMV (**Table 4, Figure 2F**).

**Table 3.** Complement analytes in patients with COVID-19 in the Yale School of Medicine cross-sectional cohort.

|  | **Non-ICU (n=14)** | **ICU (n=9)** | **p** |
| --- | --- | --- | --- |
| sC5a, pg/mL | 43.2 (43.2 – 43.2) | 77.6 (43.2 – 285.6) | 0.0016 |
| Factor D, ng/mL | 1,442 (1,234 – 1,803) | 1,825 (1,541 – 2,576) | 0.07 |

Abbreviations: ICU: intensive care unit. Statistical tests for comparison were done using the Mann Whitney U test. Values are represented as median (interquartile range).

**Table 4.**
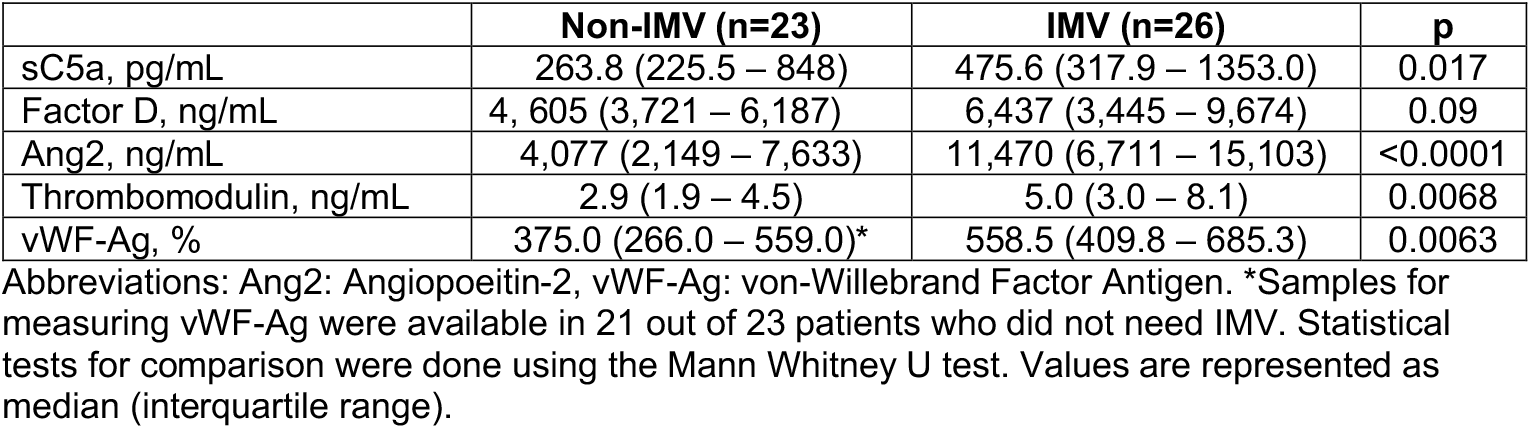
Markers of complement activation, endothelial injury and coagulation in the Yale School of Medicine longitudinal cohort.

|  | **Non-IMV (n=23)** | **IMV (n=26)** | **p** |
| --- | --- | --- | --- |
| sC5a, pg/mL | 263.8 (225.5 – 848) | 475.6 (317.9 – 1353.0) | 0.017 |
| Factor D, ng/mL | 4,605 (3,721 – 6,187) | 6,437 (3,445 – 9,674) | 0.09 |
| Ang2, ng/mL | 4,077 (2,149 – 7,633) | 11,470 (6,711 – 15,103) | <0.0001 |
| Thrombomodulin, ng/mL | 2.9 (1.9 – 4.5) | 5.0 (3.0 – 8.1) | 0.0068 |
| vWF-Ag, % | 375.0 (266.0 – 559.0)* | 558.5 (409.8 – 685.3) | 0.0063 |

Abbreviations: Ang2: Angiopoietin-2, vWF-Ag: von-Willebrand Factor Antigen. \*Samples for measuring vWF-Ag were available in 21 out of 23 patients who did not need IMV. Statistical tests for comparison were done using the Mann Whitney U test. Values are represented as median (interquartile range).

### Increase in components of the alternative pathway are associated with worse outcomes in COVID-19

We also investigated specific components of the complement cascade that may facilitate complement activation in COVID-19. In the WUSM cohort, the ratio of iC3b: C3 levels, which indicates complement activation resulting in cleavage of C3 (and is suggestive of but is not exclusively restricted to alternative pathway activation), was higher in patients needing ICU admission (**Table 2, Figure 3A**), including those requiring IMV versus those who did not (**Figure S3A**). Of note, Factor B, a component of the alternative pathway, was increased in patients with COVID-19 requiring ICU admission (**Table 2, Figure 3B**). Factor B levels also correlated with sC5b-9 levels (ρ=0.4768, **Figure 3C**). Levels of Ba, which reflect activation of the alternative pathway, were higher in patients with COVID-19 who required ICU admission (**Table 2, Figure 3D**), as well as those requiring IMV (**Figure S3B**), and those who did not survive the initial hospitalization (**Figure 3E**). The alternative pathway hemolytic activity was preserved in the COVID-19 cohort (**Table 2, Figure S3C**). Factor D was significantly higher in those who died [9,791 (4,400 – 11,579) ng/mL, n=19] compared to those who survived [4,572 (3,784 – 9,175) ng/mL, n=29, p=0.042, **Figure 3F**]. Although Factor D did not distinguish those patients requiring IMV (**Figure S3D**), it was higher in those who required renal replacement therapy [RRT, 10,158 (4432 – 12,422) ng/mL, n=9] versus those who did not [4,983 (3,786 – 10,176) ng/mL, n=39, p=0.08, **Figure S3E**]. Similar to the WUSM cohort, plasma Factor D levels of patients with COVID-19 requiring ICU admission were higher than those who did not in the Yale cohort (**Table 3, Figure S3F)**.

**Figure 3.**
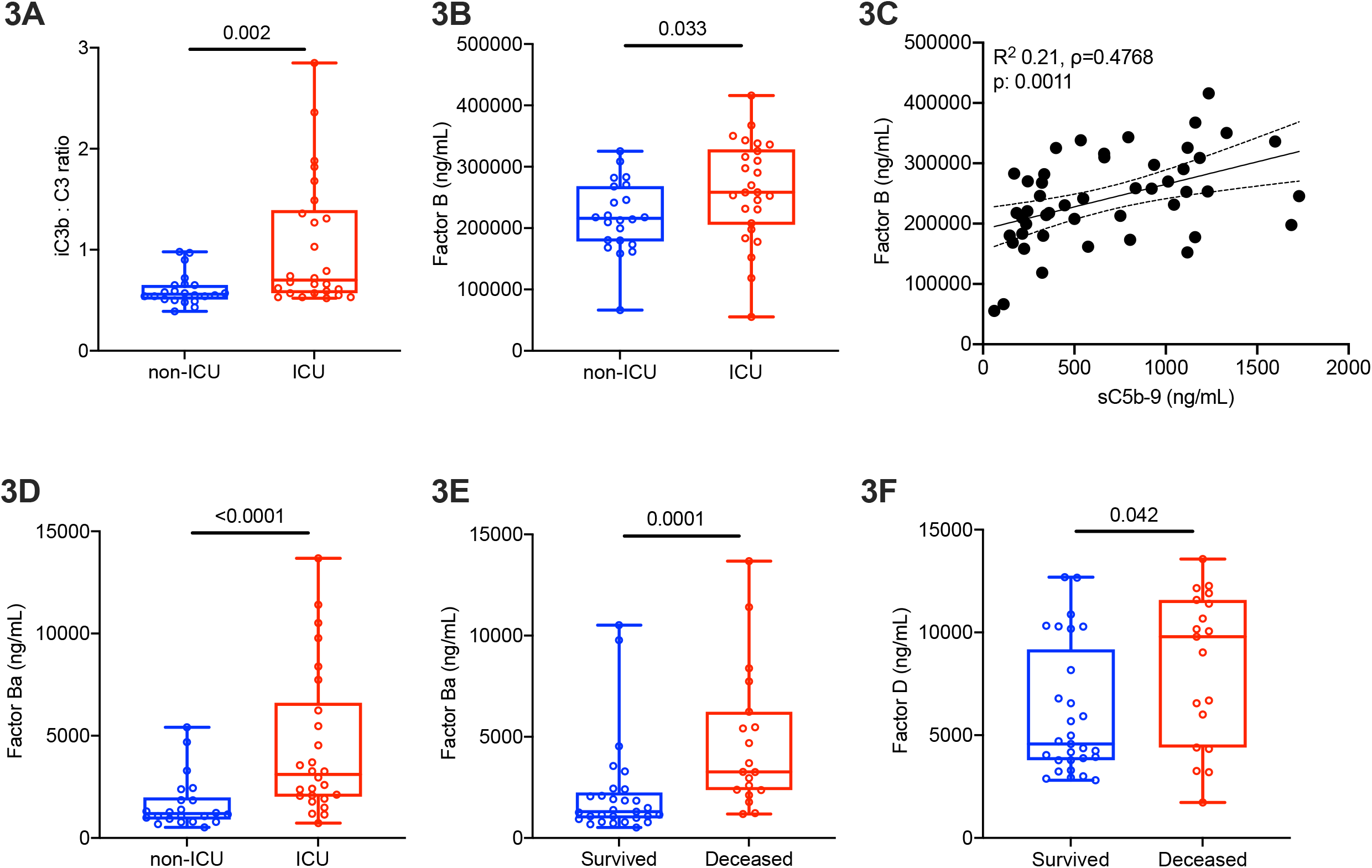
Alternative pathway activation is associated with worse outcomes in COVID-19. Comparisons in the levels of components involved in the alternative pathway (AP) in plasma of patients requiring ICU admission versus those who did not, in the WUSM COVID-19 cohort, are presented using box and whiskers plots - (A) iC3b: C3 ratio, (B) Factor B, and (D) Ba. (C) A linear regression line shows the relationship between plasma levels of sC5b-9 and Factor B. The spline chart demonstrates the mean with 95% confidence intervals. R^2^ represents the goodness-of-fit. The degree of correlation is assessed using Spearman’s Rank Correlation Coefficient test (ρ=0.4768, 95% CI 0.2146 – 0.6749, n=48). (E) Plasma Ba levels are compared in patients who survived [1,301.0 (966.0 – 2250.0), n=29] versus those who did not [3,266 (2,368 – 6236), n=19], as are the plasma levels of Factor D (F). The center of the box represents the median value, and the length of the box represents the interquartile range. The whiskers represent the minimum and maximum values in each group. Statistical significance is determined using Mann-Whitney U test.

### Complement activation is associated with markers of endothelial injury and a prothrombotic state in patients with COVID-19

Complement activation has primarily being implicated in multiorgan failure in COVID-19 due to its role in endothelial injury and inducing a prothrombotic state. Consequently, we investigated the association between Factor D and commonly utilized markers of endothelial injury, angiopoietin-2 (Ang2), and a prothrombotic state, namely thrombomodulin and the von Willebrand factor antigen (vWF-Ag). Factor D strongly correlated with Ang2 (ρ=0.5095, **Figure 4A**) and thrombomodulin (ρ=0.6050, **Figure 4B**). There was a modest correlation between Factor D and vWF-Ag (ρ=0.3367, **Figure 4C**). Ang2 was significantly higher in ICU patients with COVID-19 requiring IMV compared to those who did not (**Table 4, Figure 4D**), as was thrombomodulin (**Table 4, Figure 4E)** and vWF-Ag (**Table 4, Figure 4F)**.

**Figure 4.**
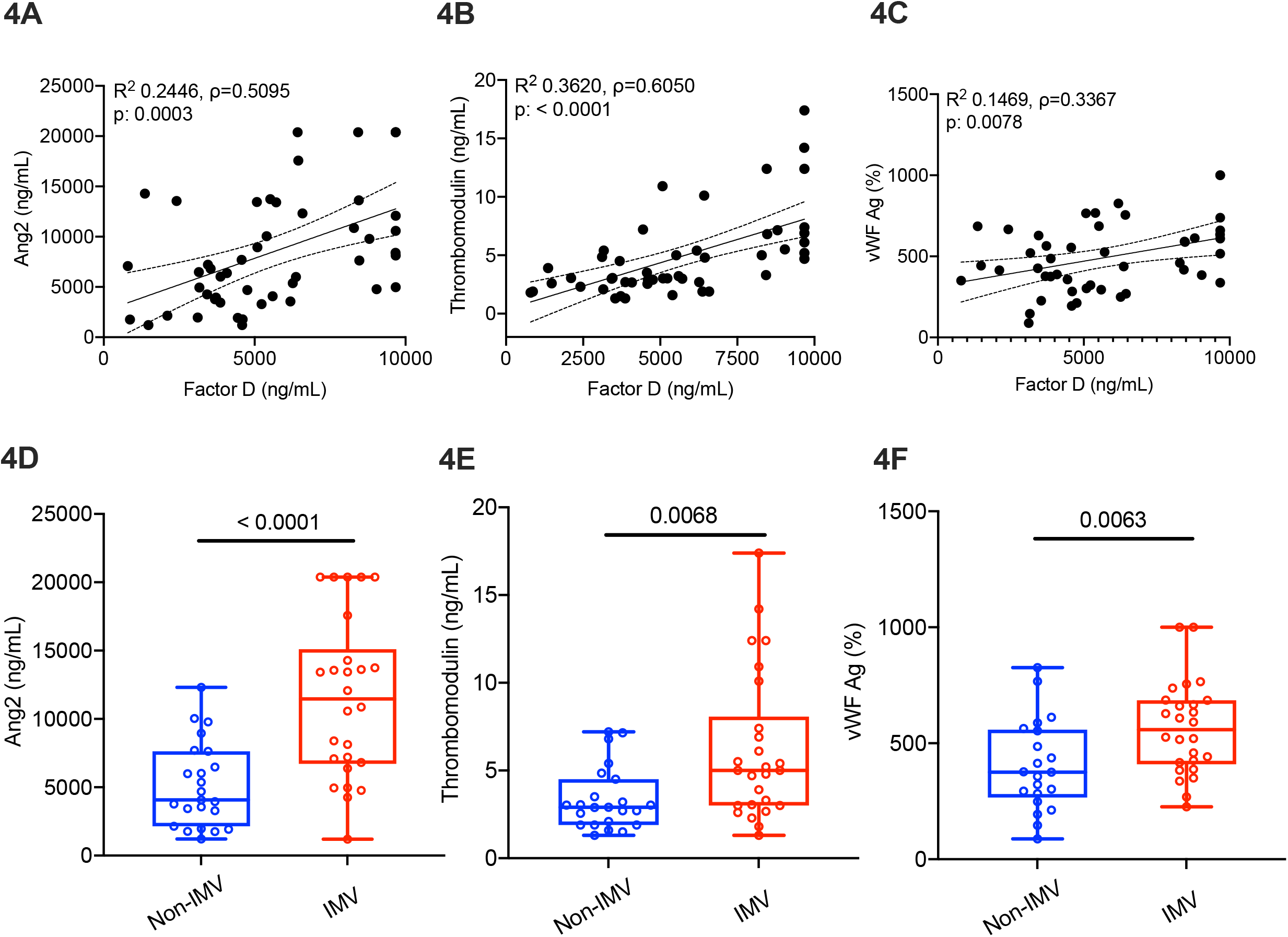
Complement activation is associated with markers of endothelial injury and a prothrombotic state in patients with COVID-19. A linear regression line shows the relationship between plasma levels of Factor D and (A) angiopoietin-2 (Ang2), (B) thrombomodulin, and (C) von Willebrand factor antigen (vWF-Ag) in the Yale longitudinal cohort (n=49). The spline chart demonstrates the mean with 95% confidence intervals. R^2^ represents the goodness-of-fit. The degree of correlation is assessed using Spearman’s Rank Correlation Coefficient test between Factor D and (a) Ang2 (ρ=0.5095, 95% CI 0.2585 – 0.6960), (b) thrombomodulin (ρ=0.6050, 95% CI 0.3829 – 0.7609), and (c) vWF-Ag (ρ=0.3367, 95% CI 0.04612 – 0.5747). Box-and-whiskers plots are utilized for comparing the levels of (D) Ang2, (E) thrombomodulin, and (F) vWF Ag in plasma of patients requiring invasive mechanical ventilation (IMV) versus those who did not. The center of the box represents the median value, and the length of the box represents the interquartile range. The whiskers represent the minimum and maximum values in each group. Statistical significance is determined using Mann-Whitney U test.

## DISCUSSION

The complement system has been implicated in COVID-19 since early in the pandemic, as evidenced by clinico-physiological and laboratory findings that supported its involvement (Java *et al*., 2020). A specific interest in COVID-19 stems from features of endothelial injury and hypercoagulability, given the cross-talk between the complement and coagulation systems (Perico *et al*., 2020). This observation has resulted in multiple phase II and III clinical trials targeting various components of the complement system (Declercq *et al*., 2020; Mastellos *et al*., 2020; Smith *et al*., 2020; Vlaar *et al*., 2020; Mansour *et al*., 2021). However, many cytokines that have been implicated in COVID-19, are also elevated in other forms of acute infection, including those leading to respiratory failure (Mudd *et al*., 2020; Bain *et al*., 2021). In certain instances, the levels of these cytokines in COVID-19 were lower than what was seen in these other diseases (Leisman *et al*., 2020; Remy *et al*., 2020). Most studies to date on the role of the complement system in COVID-19 have not included a control group of patients with another infection, or with acute respiratory failure, as a result of which it has been unclear whether complement activation is a feature of COVID-19, or is a broader indicator of critical illness. Additionally, despite multiple in vitro lines of evidence, it is unclear which specific components of the system may be associated with worse outcomes in humans with COVID-19 in vivo, which has implications for appropriately targeting this system. In this manuscript, we demonstrate that: (1) markers of complement activation are higher in severe COVID-19, compared to those hospitalized with influenza or other forms of acute respiratory failure, (2) markers of complement activation distinguish those with worse outcomes in the setting of COVID-19, in two independent cohorts, (3) the alternative pathway is activated in patients with COVID-19, and is implicated in these worse outcomes, and, (4) components of the alternative pathway associate with markers of endothelial injury and increased coagulation, which are the clinico-physiological hallmarks of severe COVID-19 vasculopathy.

We observed that markers of complement activation sare significantly higher in patients hospitalized with COVID-19, compared to those hospitalized with influenza or other forms of acute respiratory failure. Although complement activation can occur via convertase-dependent or convertase-independent pathways (e.g., thrombin cleaving C5 to C5a) in inflammatory settings such as lung injury and/or sepsis (Huber-Lang *et al*., 2006), multiple direct interactions between coronaviridae and the complement system may partly explain the elevated levels of these markers in patients with COVID-19 compared to the other etiologies. For example, the SARS-CoV spike protein can bind to mannose-binding lectin (MBL) via an N-linked glycosylation site (Zhou *et al*., 2010), initiating complement activation through the lectin pathway. The SARS-CoV-2 spike protein (subunits 1 and 2) has been shown to activate the alternative pathway in an in vitro system (Yu *et al*., 2020). Preliminary data point to the N-protein of SARS-CoV-2 mediating MASP2-drived complement activation (Gao *et al*., 2020; Kang *et al*., 2020). In comparison, the interactions between the influenza A virus (IAV) and components of the complement system appear to be more complicated. Although multiple models have demonstrated that complement activation occurs in influenza, IAV also evades complement by blocking the classical complement pathway through the M1 protein interacting with C1qA (Zhang *et al*., 2009). Additionally, microthrombi are not as common in influenza as in COVID-19, and markers of hypercoagulability appear to be higher in COVID-19 (Ackermann *et al*., 2020; Mei *et al*., 2020). These may account for some of the reasons as to why complement activation is more pronounced in COVID-19, as compared to other etiologies of acute respiratory failure, including influenza.

In two independent cohorts, we demonstrate that markers of complement activation distinguish those who had worse outcomes in the setting of SARS-COV-2 infection. The endothelial injury in COVID-19, especially in severe cases, has similarities to that seen in other forms of thrombotic microangiopathies, such as thrombotic thrombocytopenic purpura (Diorio *et al*., 2020; Java *et al*., 2020). In thrombotic microangiopathies, often a genetic predisposition, in combination with an inciting factor for endothelial damage, triggers a feed-forward loop which contributes to thrombosis and ongoing tissue injury (Java *et al*., 2020). A genetic predisposition towards complement activation has been reported in COVID-19 (Ramlall *et al*., 2020; Valenti *et al*., 2021). Additionally, infiltrating neutrophils can express prothrombotic proteins such as tissue factor (TF), both via direct expression and via neutrophil extracellular traps, driving platelet-mediated NET-driven thrombogenicity (Skendros *et al*., 2020). Infiltrating monocytes both express and release various complement components, and have anaphylatoxin receptors (i.e., C3aR, C5aR1) on their surfaces, that can bind to activated complement components (i.e., C3a, C5a). Amplifying the complement, clotting and coagulation cascades, likely contributes to severe outcomes, such as acute respiratory failure needing admission to the intensive care unit, invasive mechanical ventilation, and resulting in death, in certain cases (Carvelli *et al*., 2020).

A notable finding in our cohort, is that components of the alternative pathway are increased in COVID-19. Alternative pathway activation has been implicated in COVID-19 pathogenesis using an in vitro system (Yu *et al*., 2020). Specifically, SARS-CoV-2 spike protein has been shown to directly activate the alternative pathway, and complement-mediated killing, as well as C3c and C5b-9 deposition on TF1PIGAnull target cells was reduced by inhibition of Factor D (Yu *et al*., 2020). Transcriptomic analyses demonstrate that components of the alternative pathway (e.g., Factor B) are differentially increased in normal human bronchial epithelial (NHBE) cells infected with SARS-CoV-2 in comparison to NHBE cells infected with other respiratory infections, namely, respiratory syncytial virus, influenza (H1N1), and rhinovirus (RV16) (Blanco-Melo *et al*., 2020). Additionally, increased serum levels of Factor B have been identified using a high-throughput screen of patients with clinically severe COVID-19 (Messner *et al*., 2020). In addition to components of the complement cascade being increased in human plasma, Factor D has also been reported as being upregulated in monocytes of patients with COVID-19 pneumonia (Pekayvaz *et al*., 2021). We now show that not only is the alternative pathway activated in vivo, but Factor D also strongly correlates with markers of endothelial injury and increased coagulation in COVID-19, which are characteristic of severe disease. The next steps would be to understand the mechanistic basis for this interaction, and to evaluate whether interrupting alternative pathway-mediated activation could mitigate this vicious cycle that perpetuates tissue injury, at least in a subset of patients with severe COVID-19 who have this phenotype.

Our findings have several limitations. First, the samples were not simultaneously collected among the COVID-19, influenza, and non-COVID acute respiratory failure groups. However, they were collected in a similar time frame leading up to the SARS-CoV-2 pandemic, and subsequently processed using the same protocol, to minimize any differences in the findings. Second, we did not have levels of SARS-CoV-2 RNA to evaluate how complement activation correlates with viral load in COVID-19. A prior report would suggest that markers of complement activation do not correlate with concurrently measured viral load (Holter *et al*., 2020). This is possible, given that the patients enrolled in our study are likely presenting in the “hyperinflammatory phase” of the illness, when viral loads may be lower than the initial phase of infection (van Kampen *et al*., 2021). Third, decision-making in our ICU changed over a period of time; initially, there was a tendency for early intubation. Hence, we also included hospitalization and ICU admission in our data points; and provided data on mortality where applicable, derived from our electronic medical records. Fourth, for certain outcomes, our sample size was such that there were differences in the levels of the markers between the two groups, but they did not always meet statistical significance (e.g., mortality signal in sC5b-9); one explanation is that our study was not specifically powered for that outcome, and additionally, there may be other factors outside of acute respiratory failure that contributed to a signal such as mortality. Additionally, as we and others have previously reported, smaller differences in circulating proteins, especially in the context of complement activation, become apparent when studied locally (Daamen *et al*., 2020; Kulkarni *et al*., 2020). Reports on local complement deposition in autopsy specimens from patients with COVID-19 support this hypothesis (Bradley *et al*., 2020; Fox *et al*., 2020; Macor *et al*., 2021). We did not have adequate bronchoalveolar lavage specimens to interrogate these differences; however, this is an area of active study in our laboratory.

In summary, we show that complement activation is greater in patients hospitalized with COVID-19 when compared to those with influenza or other forms of non-COVID acute respiratory failure. Certain markers of complement activation are associated with worse outcomes, including the increased risk of ICU admission and the need for invasive mechanical ventilation in patients with COVID-19. The alternative pathway is activated in these patients, and correlates with markers of endothelial injury and increased coagulation, which are characteristics of severe COVID-19. Although our we demonstrate that increased activation of the alternative pathway is associated with worse outcomes in COVID-19, it remains to be determined whether it would be an optimal target in this disease, given the multiple mechanisms for its activation, especially in the context of acute lung injury (Irmscher *et al*., 2018; Kulkarni and Atkinson, 2020). Hence, much work remains to be done to better understand how and when to target the complement cascade, with the goal of mitigating disease severity due to SARS-CoV-2.

## MATERIALS AND METHODS

### Study Design, Settings and Participants

This study utilized plasma samples that had been independently collected from adults (aged ≥ 18 years) at two centers, Washington University School of Medicine and Yale School of Medicine.

At Washington University School of Medicine, plasma samples from patients with COVID-19 between March 26, 2020 to May 9, 2020 (‘WUSM cohort’) (Mudd *et al*., 2020). Diagnosis of COVID-19 was based on a positive nasopharyngeal swab test. Inclusion criteria required that patients be symptomatic and have a physician-ordered SARS-CoV-2 nasopharyngeal swab test performed in the course of their normal clinical care. The first available sample from the patient was utilized for analysis, primarily within 24 h of hospital admission. Other clinically relevant medical information was collected at the time of enrollment from the patient, their legally authorized representative, or the medical record.

We also report findings from influenza-infected patients enrolled in separate, ongoing studies (i.e., EDFLU study) (Turner *et al*., 2020). These patients were sampled between 2017-2020, although most were enrolled during the 2019 to 2020 influenza season, prior to the spread of COVID-19 in the St. Louis region.

To have a comparable cohort of patients with non-COVID acute respiratory failure requiring invasive mechanical ventilation, we utilized samples from the ongoing IPS (Immunity in Pneumonia and Sepsis) study at WUSM. These samples were also collected from 2019-2020 among patients admitted to the ICU, on mechanical ventilation, prior to the spread of COVID-19 in the Saint Louis region.

At Yale School of Medicine, plasma samples from 23 patients with COVID-19 were collected between April 13, 2020 to April 24, 2020 (‘Yale cross-sectional cohort’) (Goshua *et al*., 2020; Pine *et al*., 2020). A second Yale cohort (‘Yale longitudinal cohort’) was also analyzed, which included blood samples obtained longitudinally on day 1 (within 24 hours), day 4, and day 7 of hospitalization from 49 consecutive adult patients who were admitted for treatment of laboratory-confirmed COVID-19 between May 23, 2020 and May 28, 2020 and remained hospitalized until at least day 4. Diagnosis of COVID-19 was based on a positive nasopharyngeal swab test using PCR assays. Inclusion criteria required that patients be hospitalized and had a physician-ordered SARS-CoV-2 nasopharyngeal swab test performed in the course of their normal clinical care.

### Outcome Definition

Patients in the WUSM cohort were followed through May 20, 2020. Outcomes included (a) the need for ICU admission, (b) invasive mechanical ventilation (IMV), and (c) 28-day mortality. In the Yale cohorts, clinical outcomes including hospital discharge and in-hospital death were assessed. These outcomes were abstracted utilizing an honest broker system from electronic medical records.

### Sample Collection and Processing

The processing of the samples in the laboratory was similar among the cohorts. Analytes were measured in cell-free plasma collected from patients within the first 24 hours of emergency department presentation. In the WUSM COVID-19 and influenza cohorts, blood samples were collected in EDTA-containing vacutainers (BD Biosciences, San Jose, CA), transported on ice and spun down at 2,500 g (4,725 rpm) for 10 min at 4°C, after which they were stored at −80°C until further analysis (Mudd *et al*., 2020; Scozzi *et al*., 2021).

In the WUSM non-COVID (IPS) cohort, blood samples were collected in EDTA-containing vacutainers (BD Biosciences, San Jose, CA), transported on ice and spun down at 3,500 rpm for 10 min at 4°C, after which they were stored at −80°C until further analysis.

In the Yale COVID-19 cohorts, due to diurnal variations in certain analytes (e.g., plasminogen activator inhibitor-1 (PAI-1)), for hospitalized patients, blood specimens were collected with the first scheduled morning draw (i.e., occurred between 0300 h and 0700 h). For measurements of complement, coagulation and endothelial cell markers, blood was collected in 3·2% sodium citrate tubes and centrifuged at 4000 rpm at room temperature for 20 min. The resulting plasma supernatant was used for further testing. Further details on processing the samples have been reported in prior publications (Goshua *et al*., 2020; Pine *et al*., 2020).

### Measurements of complement components

#### a) Soluble C5b-9 assay

Participants were screened in duplicate for complement activation in the plasma using the soluble C5b-9 (sC5b-9) assay (BD OptEIA Human C5b-9 ELISA set, Franklin Lakes, NJ, USA) (Kulkarni *et al*., 2020). Per the manufacturer, purified native human C3, C4, C5, C6, C7, C8 and C9 were tested in the BD OptEIA™ assay at ≥ 5 mg/ml and no cross-reactivity (value ≥ 470 pg/ml) was identified.

#### b) Individual complement analytes

Individual complement analytes were evaluated in duplicate using a modified MILLIPLEX MAP Human Complement Panel 1 and 2 (MilliporeSigma, Burlington, MA, USA) based on the Luminex xMAP technology, a bead-based multiplex assay. Specifically, we employed the MILLIPLEX MAP Human Complement Panel 1 kit (HCMP1MAG) to simultaneously quantify the following analytes in the plasma: C5, C5a and Factor D. The C5 measurements are distinct from C5a, as the intact factor assays are designed such that they would not detect individual fragments based on their capture and/or detection antibodies.

Similarly, we used the MILLIPLEX MAP Human Complement Panel 2 kit (HCMP2MAG) to simultaneously quantify the following analytes in the plasma: C3, C3b/iC3b, and Factor B. Of note, the assay for C3b/iC3b detects both C3b and iC3b (per communication with manufacturer). The assay for Factor B in this kit does not detect either fragment Ba or Bb.

#### c) Alternative pathway (AP) analytes

Ba was measured in duplicate in the WUSM COVID-19 cohort using the Microvue Complement Ba fragment EIA kit (A033, Quidel Inc,San Diego, CA, USA). AP hemolysis assays were performed using a rabbit red blood cell (RBC) assay for AP activity of the plasma samples. 1 microliter of rabbit RBCs was incubated in AP buffer (gelatin veronal buffer with 20 mM MgCl_2_and 8 mM EGTA) with 10% sample concentration for 1 h. Released hemoglobin was measured at an optical density (O.D.) of 405 nm. Lysis of rabbit RBCs in water served as the positive control whereas rabbit RBCs in AP buffer served as the negative control. Hemolysis percentages were determined by an O.D. ratio using the following formula: (10% plasma with RBC in AP buffer - 10% plasma without RBC in AP buffer)/(RBC in water - RBC in AP buffer).

#### d) Measures of coagulation and endothelial cell markers

VWF antigen were measured at the Yale New-Haven Hospital (YNHH) Clinical Laboratory using ACL TOP (Instrumentation Laboratory; Bedford, MA, USA) with manufacturer reagents and controls per laboratory protocol using a latex enhanced immunoassay. The VWF antigen assay used polystyrene particles coated with rabbit polyclonal antibody directed against VWF. The results are reported as percentages compared with calibration curves using values obtained from the standardized reference population used for clinical laboratory testing throughout the YNHH system. Soluble thrombomodulin was measured using ELISA assays (Abcam, ab46508), wherein samples were diluted in a 1:4 ratio before addition to ELISA plates. Angiopoeitin-2 plasma levels were measured by Eve Technologies (Calgary, Alberta, Canada). Assays were done in duplicate according to the manufacturer’s instructions.

### Statistics

Given the sample size, we were stricter in our hypothesis testing and used non-parametric tests for comparison. Specifically, two independent groups were compared using the Mann-Whitney U test. Statistical tests for comparison were two-sided, and p < 0.05 was considered significant. In the box-and-whisker plots, the center of the box represents the median, while the length denotes the interquartile range (IQR), and the whiskers represent the minimum and maximum values in each group. The correlation between complement activation proteins and measures of endothelial injury and hypercoagulability was assessed using Spearman’s correlation, and plotted utilizing a simple linear regression line, with the error bars denoting the 95% confidence intervals. Statistical analysis was performed using IBM SPSS Statistics for Macintosh, Version 27.0 (IBM Corp, Armonk, NY), and GraphPad Prism 9 (GraphPad Software, La Jolla, CA) was employed for generating figures.

### Study Approval

The Institutional Review Board approved this study at both Washington University School of Medicine (ID#201707160, 201801209, 201808171, 201710220, 201808115, and 201910011, 201904191, 202004091, 202003085) and the Yale School of Medicine independently (IRB 2000027792 and 1401013259).

## AUTHOR CONTRIBUTIONS

HSK, JPA and HJC conceived the study; LM, SKS, JB, JM, TL, AD, DR, XW, GG, CHC, HZ, CP, PB, HR, and RS performed methodology; MC, VK, AP, MM, AMR, CWG, JAO, HSK ran software; JAO, AMR, CWG, HSK validated data; LM, MC, VK, AP, AHK and HSK performed formal analysis; LM, SKS and XW investigated; LM, MC, VK, AP, RP, AEG, CDC, AI, PM, HJC, PM curated data; HSK, JPA and HJC wrote the original draft; LM, SKS, MC, VK, JB, JM, DR, XW, AMR, CWG, JAO, RMP, AHK, AEG, CDC, and PM reviewed and edited the draft; HSK and HJC performed visualization; HSK, JPA, HJC, PM and RMP supervised, HSK administered the project, HSK, JPA, AEG, CDC, HJC, AIL, RMP and PM acquired funding.

## ACKNOWLEDGEMENTS

We thank Diane Bender and Hailey Freres from the Bursky Center for Human Immunology & Immunotherapy Programs (CHiiPs) at the Washington University School of Medicine for processing some of the samples used in our study in their core facility, Abigail Pesavento, Alexis Liszewski, Ethan Hays and Dequan Zhou for their technical assistance, and Madonna Bogacki and Lorraine Schwartz for administrative assistance.

## REFERENCES

Ackermann, M. et al.. (2020) ‘Pulmonary Vascular Endothelialitis, Thrombosis, and Angiogenesis in Covid-19’, The New England Journal of Medicine, 383(2), pp. 120–128. doi:10.1056/NEJMoa2015432.

Bain, W. et al. (2021) ‘COVID-19 versus Non-COVID ARDS: Comparison of Demographics, Physiologic Parameters, Inflammatory Biomarkers and Clinical Outcomes’, Annals of the American Thoracic Society. doi:10.1513/AnnalsATS.202008-1026OC.

Blanco-Melo, D. et al. (2020) ‘Imbalanced Host Response to SARS-CoV-2 Drives Development of COVID-19’, Cell, 181(5), pp. 1036-1045.e9. doi:10.1016/j.cell.2020.04.026.

Bradley, B. T. et al. (2020) ‘Histopathology and ultrastructural findings of fatal COVID-19 infections in Washington State: a case series’, Lancet (London, England), 396(10247), pp. 320– 332. doi:10.1016/S0140-6736(20)31305-2.

Carsana, L. et al. (2020) ‘Pulmonary post-mortem findings in a series of COVID-19 cases from northern Italy: a two-centre descriptive study’, The Lancet. Infectious Diseases, 20(10), pp. 1135–1140. doi:10.1016/S1473-3099(20)30434-5.

Carvelli, J. et al. (2020) ‘Association of COVID-19 inflammation with activation of the C5a-C5aR1 axis’, Nature, 588(7836), pp. 146–150. doi:10.1038/s41586-020-2600-6.

Cobb, N. L. et al. (2020) ‘Comparison of Clinical Features and Outcomes in Critically Ill Patients Hospitalized with COVID-19 versus Influenza’, Annals of the American Thoracic Society. doi:10.1513/AnnalsATS.202007-805OC.

Cugno, M. et al. (2020) ‘Complement activation in patients with COVID-19: A novel therapeutic target’, The Journal of Allergy and Clinical Immunology, 146(1), pp. 215–217. doi:10.1016/j.jaci.2020.05.006.

Daamen, A. R. et al. (2020) ‘Comprehensive Transcriptomic Analysis of COVID-19 Blood, Lung, and Airway’, bioRxiv, p. 2020.05.28.121889. doi:10.1101/2020.05.28.121889.

Declercq, J. et al. (2020) ‘Zilucoplan in patients with acute hypoxic respiratory failure due to COVID-19 (ZILU-COV): A structured summary of a study protocol for a randomised controlled trial’, Trials, 21(1), p. 934. doi:10.1186/s13063-020-04884-0.

Diorio, C. et al. (2020) ‘Evidence of thrombotic microangiopathy in children with SARS-CoV-2 across the spectrum of clinical presentations’, Blood Advances, 4(23), pp. 6051–6063. doi:10.1182/bloodadvances.2020003471.

Fox, S. E. et al. (2020) ‘Pulmonary and cardiac pathology in African American patients with COVID-19: an autopsy series from New Orleans’, The Lancet. Respiratory Medicine, 8(7), pp. 681–686. doi:10.1016/S2213-2600(20)30243-5.

Gao, T. et al. (2020) ‘Highly pathogenic coronavirus N protein aggravates lung injury by MASP-2-mediated complement over-activation’, medRxiv, p. 2020.03.29.20041962. doi:10.1101/2020.03.29.20041962.

Goshua, G. et al. (2020) ‘Endotheliopathy in COVID-19-associated coagulopathy: evidence from a single-centre, cross-sectional study’, The Lancet. Haematology, 7(8), pp. e575–e582. doi:10.1016/S2352-3026(20)30216-7.

Holter, J. C. et al. (2020) ‘Systemic complement activation is associated with respiratory failure in COVID-19 hospitalized patients’, Proceedings of the National Academy of Sciences of the United States of America, 117(40), pp. 25018–25025. doi:10.1073/pnas.2010540117.

Huber-Lang, M. et al. (2006) ‘Generation of C5a in the absence of C3: a new complement activation pathway’, Nature Medicine, 12(6), pp. 682–687. doi:10.1038/nm1419.

Irmscher, S. et al. (2018) ‘Kallikrein Cleaves C3 and Activates Complement’, Journal of Innate Immunity, 10(2), pp. 94–105. doi:10.1159/000484257.

Java, A. et al. (2020) ‘The complement system in COVID-19: friend and foe?’, JCI insight, 5(15). doi:10.1172/jci.insight.140711.

van Kampen, J. J. A. et al. (2021) ‘Duration and key determinants of infectious virus shedding in hospitalized patients with coronavirus disease-2019 (COVID-19)’, Nature Communications, 12(1), p. 267. doi:10.1038/s41467-020-20568-4.

Kang, S. et al. (2020) ‘A COVID-19 antibody curbs SARS-CoV-2 nucleocapsid protein-induced complement hyper-activation’, bioRxiv, p. 2020.09.10.292318. doi:10.1101/2020.09.10.292318.

Kulkarni, H. S. et al. (2018) ‘The complement system in the airway epithelium: An overlooked host defense mechanism and therapeutic target?’, The Journal of Allergy and Clinical Immunology, 141(5), pp. 1582-1586.e1. doi:10.1016/j.jaci.2017.11.046.

Kulkarni, H. S. et al. (2020) ‘Local complement activation is associated with primary graft dysfunction after lung transplantation’, JCI insight, 5(17). doi:10.1172/jci.insight.138358.

Kulkarni, H. S. and Atkinson, J. P. (2020) ‘Targeting complement activation in COVID-19’, Blood, 136(18), pp. 2000–2001. doi:10.1182/blood.2020008925.

Leisman, D. E. et al. (2020) ‘Cytokine elevation in severe and critical COVID-19: a rapid systematic review, meta-analysis, and comparison with other inflammatory syndromes’, The Lancet. Respiratory Medicine, 8(12), pp. 1233–1244. doi:10.1016/S2213-2600(20)30404-5.

Macor, P. et al. (2021) ‘Multi-organ complement deposition in COVID-19 patients’, medRxiv: The Preprint Server for Health Sciences. doi:10.1101/2021.01.07.21249116.

Magro, C. et al. (2020) ‘Complement associated microvascular injury and thrombosis in the pathogenesis of severe COVID-19 infection: A report of five cases’, Translational Research: The Journal of Laboratory and Clinical Medicine, 220, pp. 1–13. doi:10.1016/j.trsl.2020.04.007.

Mansour, E. et al. (2021) ‘Evaluation of the efficacy and safety of icatibant and C1 esterase/kallikrein inhibitor in severe COVID-19: study protocol for a three-armed randomized controlled trial’, Trials, 22(1), p. 71. doi:10.1186/s13063-021-05027-9.

Mastellos, D. C. et al. (2020) ‘Complement C3 vs C5 inhibition in severe COVID-19: early clinical findings reveal differential biological efficacy’, medRxiv, p. 2020.08.17.20174474. doi:10.1101/2020.08.17.20174474.

Mei, Y. et al. (2020) ‘Risk stratification of hospitalized COVID-19 patients through comparative studies of laboratory results with influenza’, EClinicalMedicine, 26, p. 100475. doi:10.1016/j.eclinm.2020.100475.

Messner, C. B. et al. (2020) ‘Ultra-High-Throughput Clinical Proteomics Reveals Classifiers of COVID-19 Infection’, Cell Systems, 11(1), pp. 11-24.e4. doi:10.1016/j.cels.2020.05.012.

Mudd, P. A. et al. (2020) ‘Distinct inflammatory profiles distinguish COVID-19 from influenza with limited contributions from cytokine storm’, Science Advances, 6(50). doi:10.1126/sciadv.abe3024.

Pekayvaz, K. et al. (2021) ‘Protective immune trajectories in early viral containment of non-pneumonic SARS-CoV-2 infection’, bioRxiv, p. 2021.02.03.429351. doi:10.1101/2021.02.03.429351.

Perico, L. et al. (2020) ‘Immunity, endothelial injury and complement-induced coagulopathy in COVID-19’, Nature Reviews. Nephrology. doi:10.1038/s41581-020-00357-4.

Pine, A. B. et al. (2020) ‘Circulating markers of angiogenesis and endotheliopathy in COVID-19’, Pulmonary Circulation, 10(4), p. 2045894020966547. doi:10.1177/2045894020966547.

Ramlall, V. et al. (2020) ‘Immune complement and coagulation dysfunction in adverse outcomes of SARS-CoV-2 infection’, Nature Medicine, 26(10), pp. 1609–1615. doi:10.1038/s41591-020-1021-2.

Remy, K. E. et al. (2020) ‘Severe immunosuppression and not a cytokine storm characterizes COVID-19 infections’, JCI insight, 5(17). doi:10.1172/jci.insight.140329.

Scozzi, D. et al. (2021) ‘Circulating mitochondrial DNA is an early indicator of severe illness and mortality from COVID-19’, JCI insight. doi:10.1172/jci.insight.143299.

Skendros, P. et al. (2020) ‘Complement and tissue factor-enriched neutrophil extracellular traps are key drivers in COVID-19 immunothrombosis’, The Journal of Clinical Investigation, 130(11), pp. 6151–6157. doi:10.1172/JCI141374.

Smith, K. et al. (2020) ‘A Phase 3 Open-label, Randomized, Controlled Study to Evaluate the Efficacy and Safety of Intravenously Administered Ravulizumab Compared with Best Supportive Care in Patients with COVID-19 Severe Pneumonia, Acute Lung Injury, or Acute Respiratory Distress Syndrome: A structured summary of a study protocol for a randomised controlled trial’, Trials, 21(1), p. 639. doi:10.1186/s13063-020-04548-z.

Turner, J. S. et al. (2020) ‘Impaired Cellular Immune Responses During the First Week of Severe Acute Influenza Infection’, The Journal of Infectious Diseases, 222(7), pp. 1235–1244. doi:10.1093/infdis/jiaa226.

Vabret, N. et al. (2020) ‘Immunology of COVID-19: Current State of the Science’, Immunity, 52(6), pp. 910–941. doi:10.1016/j.immuni.2020.05.002.

Valenti, L. et al. (2021) ‘Chromosome 3 cluster rs11385942 variant links complement activation with severe COVID-19’, Journal of Autoimmunity, 117, p. 102595. doi:10.1016/j.jaut.2021.102595.

Vlaar, A. P. J. et al. (2020) ‘Anti-C5a antibody IFX-1 (vilobelimab) treatment versus best supportive care for patients with severe COVID-19 (PANAMO): an exploratory, open-label, phase 2 randomised controlled trial’, The Lancet Rheumatology, 2(12), pp. e764–e773. doi:10.1016/S2665-9913(20)30341-6.

Wiersinga, W. J. et al. (2020) ‘Pathophysiology, Transmission, Diagnosis, and Treatment of Coronavirus Disease 2019 (COVID-19): A Review’, JAMA, 324(8), pp. 782–793. doi:10.1001/jama.2020.12839.

Wu, M. et al. (2020) ‘Transcriptional and proteomic insights into the host response in fatal COVID-19 cases’, Proceedings of the National Academy of Sciences of the United States of America, 117(45), pp. 28336–28343. doi:10.1073/pnas.2018030117.

Xie, Y. et al. (2020) ‘Comparative evaluation of clinical manifestations and risk of death in patients admitted to hospital with covid-19 and seasonal influenza: cohort study’, BMJ (Clinical research ed.), 371, p. m4677. doi:10.1136/bmj.m4677.

Yu, J. et al. (2020) ‘Direct activation of the alternative complement pathway by SARS-CoV-2 spike proteins is blocked by factor D inhibition’, Blood, 136(18), pp. 2080–2089. doi:10.1182/blood.2020008248.

Zhang, J. et al. (2009) ‘Influenza A virus M1 blocks the classical complement pathway through interacting with C1qA’, The Journal of General Virology, 90(Pt 11), pp. 2751–2758. doi:10.1099/vir.0.014316-0.

Zhou, T. et al. (2020) ‘Immune asynchrony in COVID-19 pathogenesis and potential immunotherapies’, The Journal of Experimental Medicine, 217(10). doi:10.1084/jem.20200674.

Zhou, Y. et al. (2010) ‘A single asparagine-linked glycosylation site of the severe acute respiratory syndrome coronavirus spike glycoprotein facilitates inhibition by mannose-binding lectin through multiple mechanisms’, Journal of Virology, 84(17), pp. 8753–8764. doi:10.1128/JVI.00554-10.

