## Supplementary Figures for "Increased complement activation is a distinctive feature of severe SARS-CoV-2 infection"

#### SUPPLEMENTARY FIGURE LEGENDS

**Figure S1. Markers of complement activation are unique to COVID-19 compared to non-COVID-19 respiratory failure.** CONSORT flow diagram showing patient enrollment, allocation and outcomes in the (A) influenza (EDFLU) cohort and (B) non-COVID acute respiratory infection (ARI) Immunity in Pneumonia and Sepsis (ISF) cohort. Box and whiskers plots of sC5b-9 levels differences between patients in the non-COVID acute respiratory failure cohort [all in ICU and requiring invasive mechanical ventilation (IMV), 243.5 (95.62 – 352.1) ng/mL], and (C) patients in the COVID-19 cohort admitted to the ICU [715.4 (448.5 – 1084.0) ng/mL], and (D) patients in the COVID-19 cohort needing IMV [922.8 (545.0 – 1198.0) ng/mL]. The center of the box represents the median value, and the length of the box represents the interquartile range. The whiskers represent the minimum and maximum values in each group. Statistical significance is determined using Mann-Whitney U test.

**Figure S2. Complement activation is associated with worse outcomes in patients with severe COVID-19.** Box and whiskers plots of plasma sC5b-9 levels in the WUSM COVID-19 cohort (A) in patients needing to be hospitalized versus those discharged (not hospitalized) from the emergency room, and when the COVID-19 cohort was restricted to (B) hospitalized patients needing invasive mechanical ventilation [IMV, 922.8 (545.0 – 1198.0) ng/mL, n=29] versus those who did not [620.8 (399.9 – 851) ng/mL, n=95], and (C) patients admitted to the ICU who needed IMV [922.8 (545.0 – 1198.0) ng/mL, n=29, same as in S2B] versus those who did not [623.3 (352.5 – 936.6) ng/mL, n=43]. Comparison of plasma sC5a levels among those patients who needed ICU admission versus those who did not in (D) the WUSM cohort, and (E) the Yale cross-sectional cohort. The center of the box represents the median value, and the length of the box represents the interquartile range. The whiskers represent the minimum and maximum values in each group. Statistical significance is determined using Mann-Whitney U test.

**Figure S3. Components of the alternative pathway are associated with worse outcomes in COVID-19.** Box and whiskers plots of (A) plasma iC3b : C3 ratio of patients from the WUSM COVID-19 cohort requiring invasive mechanical ventilation [IMV, 0.72 (0.58 – 1.38), n=17] versus those who did not [0.56 (0.52 – 0.66), n=28], and (B) plasma Ba levels of patients from the WUSM COVID-19 cohort requiring IMV [2,961 (2,001 – 8,065) ng/mL] versus those who did not [1,386 (1,072 – 3,262) ng/mL]. (C) Alternative pathway hemolytic activity in the WUSM COVID-19 cohort, expressed as a percentage. Comparison of plasma factor D levels in patients from the WUSM COVID-19 cohort who (D) required IMV [6,690 (4,286 – 11,036) ng/mL] versus those who did not [5,920 (3,782 – 10,283) ng/mL] and (E) required renal replacement therapy (RRT) versus those who did not. (F) Factor D levels in patients from the Yale cohort who needed ICU admission versus those who did not. The center of the box represents the median value, and the length of the box represents the interquartile range. The whiskers represent the minimum and maximum values in each group. Statistical significance is determined using Mann-Whitney U test.

### Figure S1

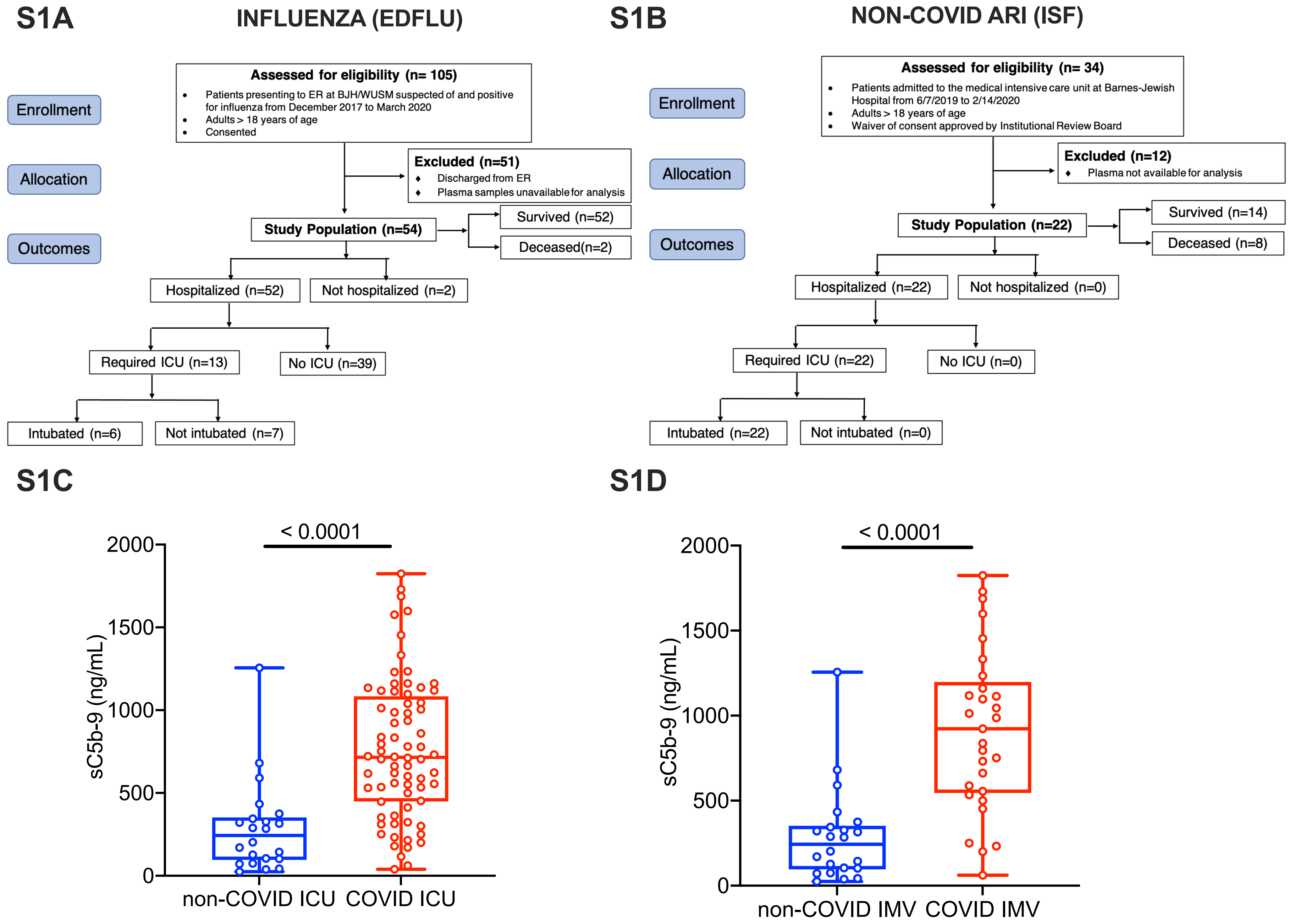
S1A
INFLUENZA (EDFLU)
Enrollment
Allocation
Outcomes
Assessed for eligibility (n= 105)
Patients presenting to ER at BJH/WUSM suspected of and positive for influenza from December 2017 to March 2020
Adults > 18 years of age
Consented
Excluded (n=51)
Discharged from ER
Plasma samples unavailable for analysis
Study Population (n=54)
Survived (n=52)
Deceased(n=2)
Hospitalized (n=52)
Not hospitalized (n=2)
Required ICU (n=13)
No ICU (n=39)
Intubated (n=6)
Not intubated (n=7)
S1B
NON-COVID ARI (ISF)
Enrollment
Allocation
Outcomes
Assessed for eligibility (n= 34)
Patients admitted to the medical intensive care unit at Barnes-Jewish Hospital from 6/7/2019 to 2/14/2020
Adults > 18 years of age
Waiver of consent approved by Institutional Review Board
Excluded (n=12)
Plasma not available for analysis
Study Population (n=22)
Survived (n=14)
Deceased (n=8)
Hospitalized (n=22)
Not hospitalized (n=0)
Required ICU (n=22)
No ICU (n=0)
Intubated (n=22)
Not intubated (n=0)
S1C
< 0.0001
sC5b-9 (ng/mL)
2000
1500
1000
500
0
non-COVID ICU
COVID ICU
S1D
< 0.0001
sC5b-9 (ng/mL)
2000
1500
1000
500
0
non-COVID IMV
COVID IMV

### Figure S2

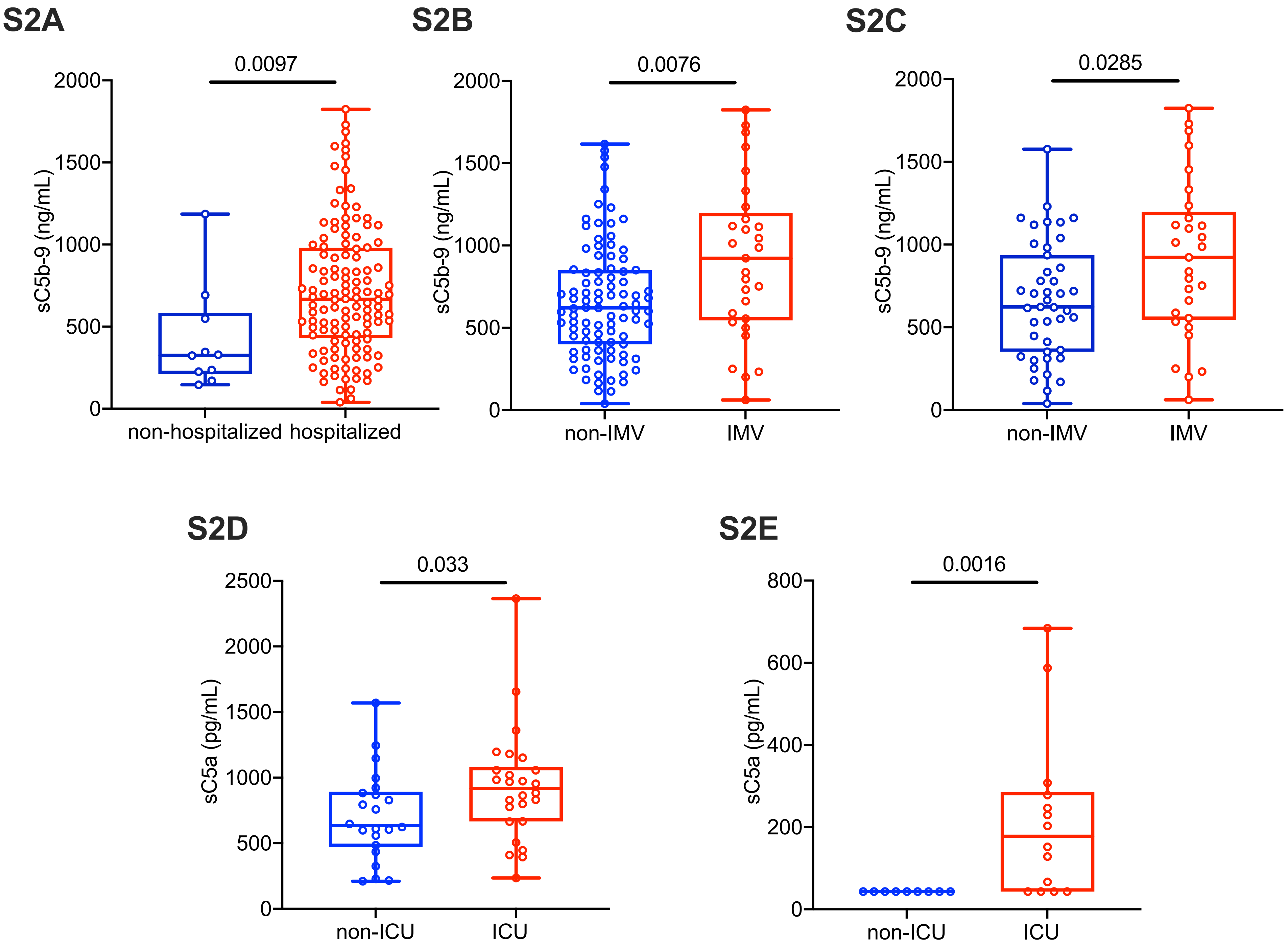
S2A
0.0097
2000
1500
1000
500
0
sC5b-9 (ng/mL)
non-hospitalized
hospitalized
S2B
0.0076
2000
1500
1000
500
0
sC5b-9 (ng/mL)
non-IMV
IMV
S2C
0.0285
2000
1500
1000
500
0
sC5b-9 (ng/mL)
non-IMV
IMV
S2D
0.033
2500
2000
1500
1000
500
0
sC5a (pg/mL)
non-ICU
ICU
S2E
0.0016
800
600
400
200
0
sC5a (pg/mL)
non-ICU
ICU

### Figure S3

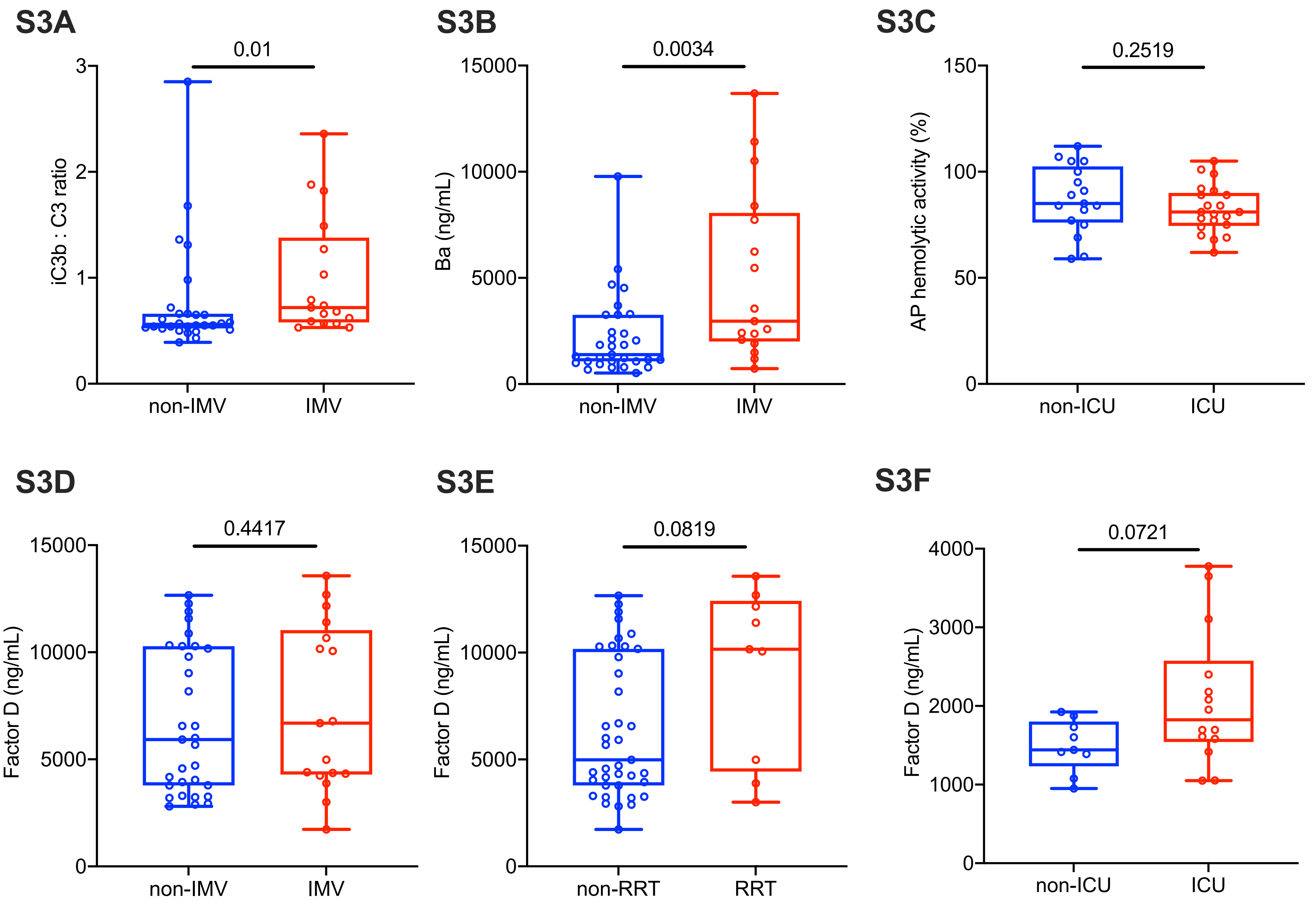
S3A
0.01
iC3b : C3 ratio
3
2
1
0
non-IMV
IMV
S3B
0.0034
Ba (ng/mL)
15000
10000
5000
0
non-IMV
IMV
S3C
0.2519
AP hemolytic activity (%)
150
100
50
0
non-ICU
ICU
S3D
0.4417
Factor D (ng/mL)
15000
10000
5000
0
non-IMV
IMV
S3E
0.0819
Factor D (ng/mL)
15000
10000
5000
0
non-RRT
RRT
S3F
0.0721
Factor D (ng/mL)
4000
3000
2000
1000
0
non-ICU
ICU
